# Habitat restoration promotes recolonisation by extirpated species in model meta food webs

**DOI:** 10.64898/2026.06.15.731902

**Authors:** Lucie Thompson, Miguel Lurgi

## Abstract

Successful ecosystem restoration is intimately linked to the persistence of species in local communities and across landscapes. As such quantitative approaches to ecological restoration require the integration of community and metapopulation ecology. Together these disciplines demonstrate that local colonisation, via habitat connectivity and size, and species interactions, both modulate the process of community assembly. However, thus far restoration ecology still remains disconnected from network ecology this preventing a holistic, community-wide perspective to restoration.

We aim to inform ecological restoration using a multi-layer modelling framework integrating ecological interactions and species’ dispersal dynamics. We explore the drivers that modulate recolonisation dynamics of species across restored landscapes. We further investigate how restoration improves the structural properties of food webs, the number of successful recolonisations and the role of configuration of restored patches in restoration outcomes.

We find that recolonisation is the result of a trade-off between dispersal ability and energy requirements. 97% of plant recolonisation and 88% of herbivore recolonisations happened within close proximity to the source patches. Better dispersers – intermediate and top species in the food webs – were able to recolonise habitat by benefitting from the increased biomass influx from restoration. When only a small proportion of the landscape could be restored, the location and connectivity of restored areas strongly influenced the outcome of restoration: more connected patches enabled on average the recolonisation of about 1 additional intermediate species compared to that of isolated patches. However, this difference faded as soon as more patches were restored, and improving larger portions of the landscape always resulted in better outcomes. Restoring 1/3 of the landscape enabled on average the recolonisation of ~4 additional species. Our findings suggest that quantitative models can inform restoration efforts necessary to bring native species back to restored areas. They also suggest that attention should be given to the requirements of the recolonisers, the distance of their introduction from restored areas and their trophic and ecological niche. These aspects are crucial to assess their energy and habitat requirements for successful establishment.

## Introduction

Through passive methods such as the abandonment of an environmentally degrading activity or active methods such as landscape remodelling, habitat enhancement by tree planting or the input of nutrients, restoration ecology aims at “[…] assisting the recovery of an ecosystem that has been degraded, damaged or destroyed” (Society for Ecological Restoration International, 2004; Gann *et al*., 2019). Restoration projects can span from freshwater (Bernhardt *et al*., 2005), to forests (Stanturf *et al*., 2014), grassland (Török *et al*., 2021) or dryland systems (Shackelford *et al*., 2021) and can have varied objectives: from restoring community structure, to bringing back native species or mitigating the effects of climate change (Stanturf *et al*., 2014). Given their very wide applications, it is clear that the task of developing theory that offers predictability across such different objectives and systems is difficult, but increasingly needed (Palmer *et al*., 2016). Realistic multi-layer studies that integrate ecological processes operating across scales (e.g. species dispersal) and ecological interactions (e.g. food webs) influencing population dynamics at local scales (Urban *et al*., 2016) could be an interesting tool to inform restoration projects

Restoration ecology draws on knowledge from many disciplines (Gann *et al*., 2019), overlapping with a lot of “classical” ecological theory (Palmer *et al*., 2016). To explore how these multi-layer studies could inform restoration ecology, efforts at developing predictive theory should incorporate at least three central aspects of ecological research. First, food web ecology - how trophic interactions redistribute energy across ecosystems (Lindeman, 1942; Elton, 1958) – can inform outcomes of recolonisation based on their effects on entire communities (Vander Zanden *et al*., 2006). Second, community and invasion ecology focus on aspects of ecological successions, the role of initial conditions and recoloniser traits in modulating recolonisation. Lastly, metapopulation ecology allows us to consider the fact that restored patches are embedded within a larger landscape and integrate spatial influence of surrounding sites on restored sites and vice-versa (Maurer, 2006).

Numerous examples illustrate the importance of integrating ecological interactions (such as trophic relationships) into studies aimed at understanding recolonisation dynamics and their sometimes unintended consequences (Vander Zanden *et al*., 2006). The return of wolves (*Canis lupus*) to Yellowstone National Park in the 1990s, for example, indirectly allowed the recovery of overgrazed riparian vegetation by elks (*Cervus elaphus*), and prompted an increase in the populations of beaver (*Castor canadensis*) and bison (*Bison bison*) (Beschta & Ripple, 2009). More generally, food web dynamics can provide insights into how species (re)introductions would be expected to restructure ecosystems from the top down. Similarly, from the bottom up, food web theory explains how ecosystem productivity is related to food chain length (Pimm, 1982) through the energy limitation hypothesis (Lindeman, 1942; Elton, 1958). These concepts give insights into how to encourage the recolonisation of top predators into disturbed ecosystems and improve key ecosystem properties such as food chain length. In spite of this compelling evidence pointing at the role of species interactions in restored ecosystems dynamics as well as recent studies (Bellmore *et al*., 2017; Loch *et al*., 2020) advocating for the incorporation of food webs into frameworks aimed at informing restoration projects, they have been mostly overlooked in favour of simpler, single species-centred approaches (Zanden *et al*., 2016; Bellmore *et al*., 2017).

Similarly in restoration, there is ample evidence that initial conditions and species composition can modulate recolonisation via priority effects (Palmer *et al*., 1997; Weidlich *et al*., 2021), ultimately impacting the speed and magnitude of post-restoration recovery (Manhães *et al*., 2022). Although landscape recolonisation in the context of restoration, i.e. by native species previously displaced from an area due to habitat degradation or harvesting, might follow different dynamics than the invasion of exotic species, this difference should mainly lie in their effect on established communities. Thus, lessons on recolonisation dynamics can likely be learned from studies integrating trophic interactions to explore invasions of food webs by exotic species (Romanuk *et al*., 2009; Lurgi *et al*., 2014; Häussler *et al*., 2021; Sentis *et al*., 2021). Of particular relevance is the importance of food web structure (Elton, 1958) and invaders’ traits in modulating invasion success. For example, food web connectance and species diversity, alongside trophic level of the invasive species are known to interact and modulate invasion success (Frost *et al*., 2019).

Restored patches of habitat can only benefit from species recolonisation if they are not isolated but embedded within landscapes that enable the exchange of individuals with surrounding habitats (Brown & Kodric-Brown, 1977; Gravel *et al*., 2010). Thus, frameworks that integrate spatial processes with local species interactions are particularly relevant for exploring restoration scenarios and recolonisation events. In the context of invasions, Häussler *et al*. (2021) using theoretical models of meta food webs - collections of patches that harbour trophic networks and exchange individuals through dispersal - found that invasion increased with nutrient supply but that aspects of landscape connectivity could halt the spread of invasive species (see also Buxton *et al*., 2014 for an empirical example). More generally, the integration of dispersal and connectivity processes can give rise to interesting dynamics whereby landscape heterogeneity and notably habitat restoration can promote diversity and stability through source-sink dynamics (Gravel *et al*., 2010) driven by rescue (Brown & Kodric-Brown, 1977) and drainage effects (Ryser *et al*., 2021). This suggest that restoration efforts in combination with effective landscape connectivity should allow for the recolonisation of extirpated species.

Overall, knowledge on landscape connectivity, ecosystem productivity, and trophic interactions of ecological relevant species could help better predict restoration outcomes for community reassembly in the context of habitat restoration. We explore this question here using meta food web models. Formally, meta food webs, a special case of meta communities (Holyoak *et al*., 2005), are defined as a collection of spatially connected food webs. Local food webs, or “patches,” behave according to classical population dynamics models; specifically in our study, complex bioenergetic food webs comprising plants, intermediate species, and top predators. Using this framework we simulated targeted restorations (increase in patch quality) on initially “degraded” landscapes. We monitored food web structure and recolonisation events before and after restoration to understand (1) how restoration (via habitat improvement and species recolonisations) affects food webs across the landscape, (2) which pre-restoration food web and landscape features best predict these recolonisation events. We also tested different restoration scenarios by varying the position of restored patches throughout the landscape in either a clustered or scattered manner to explore (3) how different clustering scenarios of restored patches affects recolonisation dynamics.

## Methods

We modelled bio-energetic meta food webs using a modified version of the allometric model proposed by Ryser *et al*. (2019, 2021) and Häussler et al. (2021), where feeding (i.e. trophic) interactions between species are determined by the niche model (Williams & Martinez, 2000) and species metabolism, consumption and dispersal are dependent on body size. The niche model is a simple model that generates networks with key structural properties observed in empirical food webs by fixing two parameters, the number of species and fraction of realised interactions (i.e., the connectance) of the network. From these feeding links species’ body masses are calculated based on their trophic positions following Williams & Martinez (2004). Body mass in turn determines consumers’ dispersal capacities and metabolic rates (see model specifications below) (Häussler *et al*., 2021; Ryser *et al*., 2021). Population dynamics in this model are derived using a set of coupled ordinary differential equations, where the rate of change of species’ biomass is determined by the sum of biomass gains and losses through feeding (for consumers) or intrinsic growth (for basal resources), predation, metabolism (death rate), emigration, and immigration.

We generated 6 different landscapes comprising 15 habitat patches randomly placed on a square-shaped space and harbouring local independent food webs. All patches initially shared the same abiotic conditions (i.e. growth rates of basal species do not vary across patches), and our model does not consider nutrient dynamics explicitly (contrary to Häussler *et al*., 2021 and Ryser *et al*., 2021 who included nutrient refreshment rates). On this spatial food web setting, we ran numerical simulations of species populations dynamics. After letting initial dynamics stabilise, we simulated restoration over the initial degraded scenario by increasing the growth rates of basal resources three-fold on a fraction of patches in the landscape, from 1 up to 5 restored patches. Increases of growth rates in basal species (e.g. plants) are meant to represent increased productivity within the habitat patch. We tested two configurations of landscape restoration, one where restored patches were clustered and one where they were scattered randomly across the landscape. Concomitantly to restoration, we let any species from the regional pool that did not survive in the initial low quality (i.e. degraded) landscape attempt to recolonise the system through the patch closer to the top-left corner of the landscape (Figure 1; inspired by Häussler *et al*., 2021). This effectively simulates the re-colonisation of restored landscapes by species that were previously excluded due to the low quality of the habitat (an effect of human disturbance on natural landscapes).

**Figure 1.**
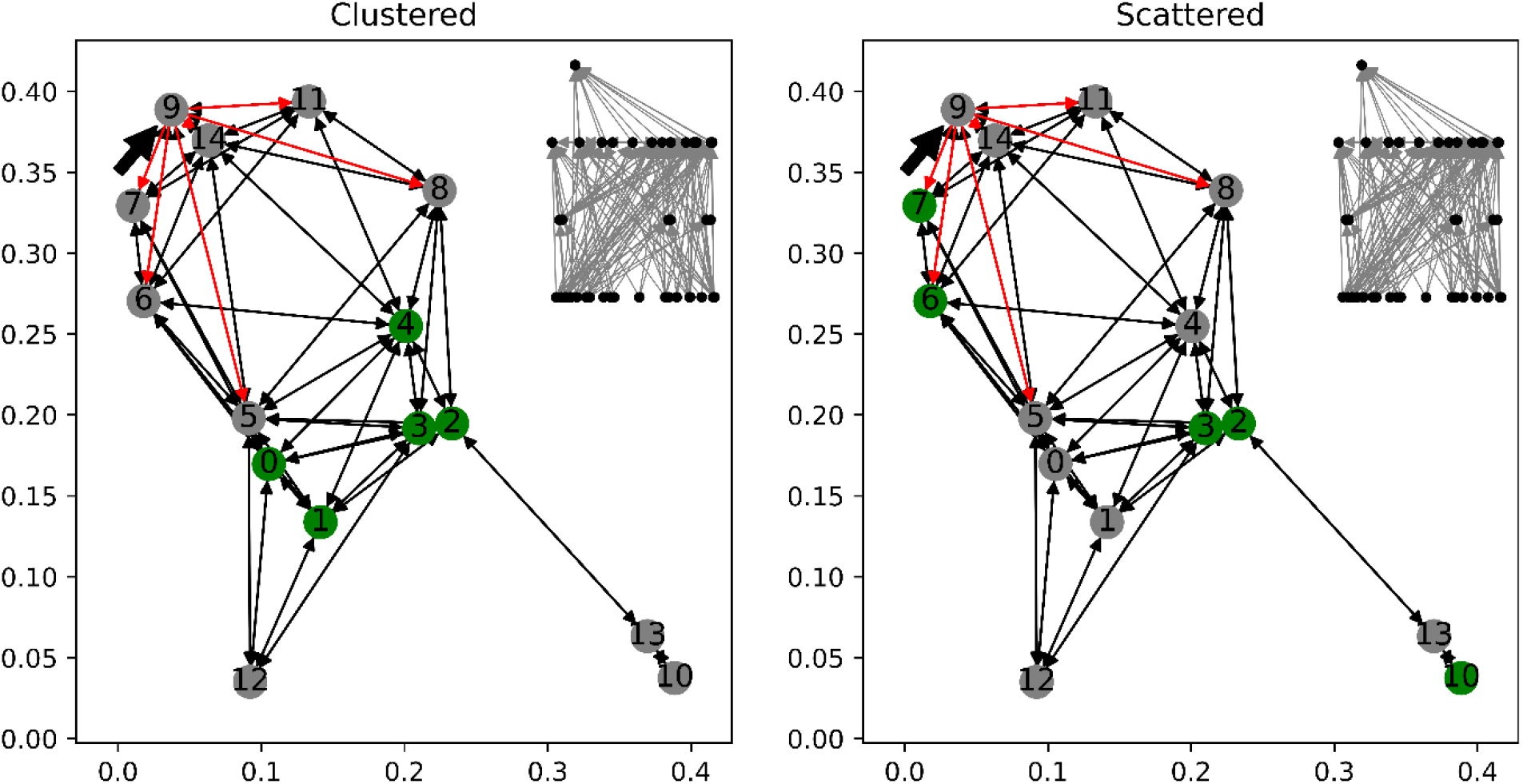
Illustration of a landscape under either a clustered (left) or scattered (right) patch restoration scenario. Map of one of our 5 randomly generated landscape with 15 patches coloured by their restoration status (grey = not restored - patches were kept with the same “low” habitat quality Δ*r*_*z*_ as pre-restoration, green = restored - the growth rate of plants on those patches was increased threefold). Left panel: clustered scenario where restored patches were clustered in a circle of radius 0.05 centred around the middle of the landscape. Right panel: scattered scenario, where restored patches and restoration sequence were randomly drawn from uniform distributions. Arrows between nodes in the network of patches illustrate the connectivity of patches from the perspective of a median disperser with maximal dispersal distance (eq. 6) *δ*_*median*_ = 0.2, which corresponds to the median dispersal distance among species from the regional food web. The top left thick arrow points to the source patch that was used to re-introduce recolonising species. Red arrows leaving the source patch illustrate the dispersal paths to patches that can be reached in one single dispersal event from it by a median disperser. The network in the top right corner of the plots represents an example of a realised food web that might occur in a given patch (here patch “8”). Nodes in the food webs (black points) represent species and edges (arrows) represent trophic links. Arrows point towards the predator and represent biomass flux. Species are organised by trophic level.

### Landscape generation

Landscapes consisted of randomly distributed patches on a lattice of 0.4 by 0.4 arbitrary units. We ensured that patches were separated by at least 0.01 units by redrawing their coordinates when this condition was not met. To compare restoration scenarios of clustered versus scattered patches, where clustered patches were situated in the middle of the landscape, we enforced that the 5 middle patches be contained within a circle of radius 0.05 centred around the middle of the landscape. The 10 remaining patches were randomly placed outside this circle. The dimensions of the landscapes were chosen to ensure that species’ median dispersal range *δ*_*i*_ (equation 5 below) across all species in the food web is close to the median inter-patch distance, ensuring that at least half of the species in the communities can effectively disperse across the landscape. Thus, all patches were potentially connected to all others, but accessible to different extents by different species depending on their dispersal capacities (see equation 4 below).

### Food web dynamics

We generated 5 distinct regional food webs - a regional pool of species and all their potential feeding interactions - comprising 100 species and with connectance = 0.1 using the niche model (Williams & Martinez, 2000). We selected those 5 regional food webs ensuring that they could host at least 55 species in their isolated state (i.e. when not in the metacommunity). This was achieved by running their dynamics under low habitat quality (see below) and keeping those with at least 55 persisting species after dynamics had stabilised (see below for stability criteria).

As in Häussler *et al*. (2021) and Ryser *et al*. (2021), local (patch-level) population dynamics across patches were modelled using a bio-energetic model of species interactions. In this model, in a system with *S* species, the rate of change of consumer *i*’s biomass density on patch *z B*_*i,z*_ (Eq. 1), depends on the feeding rate *F*_*i,j*_ of species *i* on species *j*, the metabolic rate *x*_*i*_ of species *i* and dispersal dynamics encompassed in the emigration and immigration terms *E*_*i,z*_ and *I*_*i,z*_.

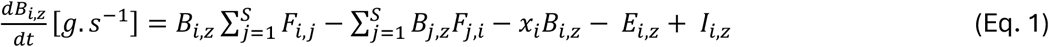

Similarly, the rate of change of basal resource species’ (e.g. plants) biomass density *P*_*i,z*_ (Eq. 2) depends on their intrinsic growth rate *r*_*i,z*_, modulated by a patch quality term Δ*r*_*z*_ that improves the growth rate of the species on patch *z* in scenarios of enhanced habitat quality.

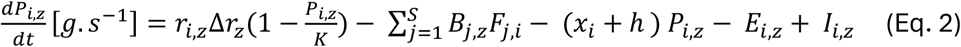

In what follows, “landscape heterogeneity” refers to differences in basal resources growth rates across patches in the landscape by exploring different values of Δ*r*_*z*_. By multiplying *r*_*i,z*_ by a patch-specific constant Δ*r*_*z*_ we created better- or worse-quality patches, thus varying landscape heterogeneity. “Low quality” patches had Δ*r*_*z*_ = 0.5 whilst “high quality” patches had Δ*r*_*z*_ = 1.5.

### Body mass scaling of vital rates

Species’ body mass (Eq. 3) was inferred from their trophic position in the food web according to Eq. 3. A minimum arbitrary body mass *m*_0_ of 0.01 g was set for resource species. Their carrying capacity *K* = 1.3771 *g*^−1^ was constant and calculated from 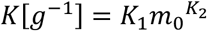 with *K*_1_ = 5 and *K*_2_ = 0.28 (Sentis *et al*., 2021). All other species’ body masses *m*_*i*_ scaled with their prey-averaged trophic level *TP*_*i*_ (Williams & Martinez, 2004) assuming an average predator-prey body mass ratio 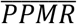 of 100.

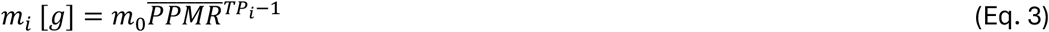

Net dispersal of species comprises the sum of both emigration and immigration terms, which occurred concomitantly with other drivers of population dynamics. Emigration, or the rate at which a species leaves a given patch, depends on the per capita emigration rate *d*_0_, constant and set to 1*e*^−8^ *s*^−1^ (Eq. 4).

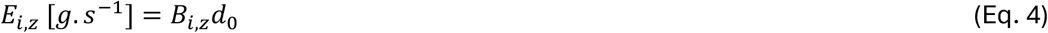

Immigration into a given patch, on the other hand, corresponded to the sum of immigrants from all neighbouring patches that succeeded to reach that patch. Immigration rates of species *i* to patch *z* (Eq. 5) depend on species *i*’s success rate when dispersing from patch *n* to *z*, (1 − *δ*_*i,nz*_), where *δ*_*i,nz*_ is the ratio of the distance between patches *n* and *z* relative to species *i*’s maximum dispersal range *δ*_*i*_ (Eq. 6) such that 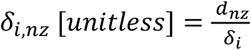.

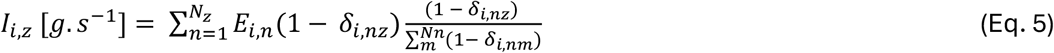

Consumers’ maximum dispersal range *δ*_*i*_ scaled with their body mass *m*_*i*_, with a slope ε = 0.05 and intercept *δ*_0_ = 0.1256 (Eq. 6; Häussler *et al*., 2021). Thus, larger species were more mobile and able to disperse further.

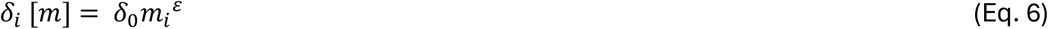

Resources’ maximal dispersal ranges were drawn from a uniform probability distribution between 0 and 0.5 (Häussler *et al*., 2021).

The fraction 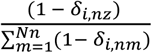 in Eq.5 accounts for the fraction of species *i*’s biomass that travelled towards patch *z* from patch *n* relative to that of all its *Nn* other neighbouring patches also accessible to species *i*. This formulation reflects the realistic assumption that a larger fraction of biomass disperses to closer patches.

Per unit biomass feeding rate of species *i* on species *j* (Eq. 7) was defined as a Holling type II function.

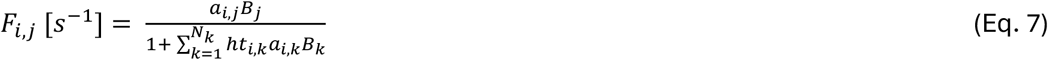

Both the handling time *ht*_*i,j*_ (Eq.8) and the attack rate *a*_*i,j*_ (Eq. 9) of species *i* on species *j* (rate of successful attacks, that accounts for the search area a species can cover) scaled with body size of both consumer and resource species *m*_*i*_ and *m*_*j*_.

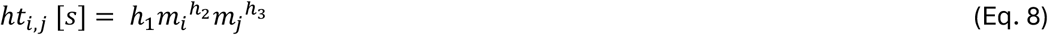

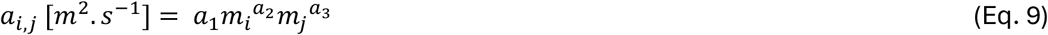

Parameter values for the scaling of attack rate and handling times with body mass were taken from (Binzer *et al*., 2016), originally derived from (Rall *et al*., 2012) and references therein. In particular, the intercept *a*_1_ in the attack rate was set to *a*_1_ = *e*^−13.1^, while the scaling coefficient for consumer species *a*_2_ = −0.8 and the scaling coefficient for resource species *a*_3_ = 0.25 (Rall *et al*., 2012). Similarly, the intercept *h*_1_ for the handling time was set to *e*^9.66^, whereas the scaling coefficient for consumer species *h*_2_ = 0.47 and the scaling coefficient for resource species *h*_3_ = −0.45 (Rall *et al*., 2012).

Consumers’ metabolic rate (death rate) *x*_*i*_ also scaled with body size according to Eq. 10:

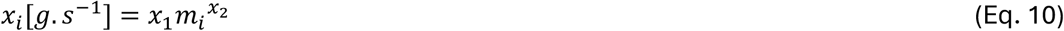

with *x*_1_ = *e*^−16.54^ and *x*_2_ = −0.31 (Rall *et al*., 2012).

### Simulation protocol

The model described above was simulated numerically to obtain population dynamics of species across patches. We performed a series of replicated experiments exploring different scenarios of ecosystem restoration. In-silico experiments were all run on five landscapes and five independently drawn regional food webs. The following protocol was applied: (1) Initial population dynamics were allowed to stabilise on a homogeneous low-quality landscape (all patches had Δ*r*_*z*_ = 0.5). Then (2) re-colonisation was allowed in the homogeneous landscape with no restored patches. Finally, (3) restoration (where Δ*r*_*z*_ = 1.5) was simulated independently on 1 to 5 patches while allowing recolonisation. Thus, each restoration replicate involved a fixed number of restored patches (as opposed to cumulative restoration in several “stages”).

Throughout the simulations, species were considered extinct when their biomasses went below a threshold of 1e^-8^. To avoid numerical problems with species’ biomasses becoming increasingly smaller, simulations were halted when an extinction happened, biomasses of extinct species were “manually” set to zero and simulations were then restarted from the state of all the other species before halting. Simulations were stopped when demographic measures remained approximatively constant with time (i.e. approaching a steady state). Concretely, we sub-divided the last 25% of the simulation timesteps into five non-overlapping time-windows. We then computed species’ mean biomass within each time-window. We assessed the coefficient of variation of these mean biomasses in time across the five time-windows. If coefficients of variation of all species’ biomasses across the five time-windows were smaller than 1e^-2^, the dynamics were considered to have stabilised, and the simulation was halted. This allowed for both oscillating and non-oscillating systems to be considered ‘stable’ if they were close enough to a steady state. If after 12 hours of runtime, demographic measures didn’t meet our stable state criteria then they were stopped and discarded from further analyses.

#### Initial population dynamics

Each patch in the landscapes was initialised with a random sample of 33 species from the regional pool of 100 species. A random seed was used to ensure that landscape replicates for the same food web were initialised with the same regional species pool so that potential differences in community compositions between landscapes would emerge from differences in their spatial configuration only. All patches were started with the same abiotic conditions (i.e. Δ*r*_*z*_ = 0.5 across all patches). Initial biomass of resources was set to their carrying capacity (see *Body mass scaling of vital rates* above) and initial biomass of consumers to 1/8 of that value.

#### In-silico recolonisation experiments

Once initial dynamics had stabilised, we allowed for recolonisation by species from the regional pool that were initially unable to persist on the recolonisation patch (top-left patch; Figure 1 – patch with arrow) or that were not randomly seeded on that patch in the initial landscape. This was achieved by initialising the biomass of all species absent in the recolonisation patch but present in the regional pool to 1/100th of the abundance of extant species in the patch.

Before starting restoration, and to assess the effects of recolonisation alone, independently from restoration, we first allowed for recolonisation of the homogeneous, pre-restoration (i.e. degraded) landscape. This recolonised homogeneous landscape represents our “zero patches improved” control throughout the study.

#### Restoration experiment

To assess the effect of habitat restoration on complex meta food webs, we simultaneously improved a set number of patches from 1 to 5 (independent simulations were conducted for each number of improved patches) by increasing their Δ*r*_*z*_ threefold, from 0.5 to 1.5 on all our combinations of landscapes (5) and food webs (5). This protocol resulted in six improvement levels with 0 to 5 patches improved, testing how restoring an ever increasing fraction of the landscape impacted local communities’ composition and dynamics. For example, for the restoration of three patches in the landscape we would (1) initialise species’ biomasses to the pre-restoration and pre-recolonisation equilibrium, (2) simultaneously improve three patches and (3) initialise recolonisers from the regional pool with a positive biomass in the top-left patch (see *In silico recolonisation experiments* above).

Two restoration configurations were tested: clustered and scattered (Figure 1). In the clustered configuration, the five improved patches were the five most geographically central patches in the landscape (see *Landscape generation* section above). For the scattered configuration, we generated five different and independent sequences of five patches drawn randomly without replacement from the pool of patches. These sequences determined which patches were to be restored and the order of restoration. Thus, for example, for a randomly drawn restoration sequence [2, 5, 7, 12, 6], the first patch improved would be patch number 2, then patches [2, 5] simultaneously, then [2, 5, 7], and so on until the whole set of 5 chosen patches are restored. Sequences were kept constant across simulations. We tested the effect of restoration sequence on restoration outcome by assigning each sequence an ID (see Evaluating the relative importance of initial conditions and landscape properties section below).

Patch restoration was always conducted from the initial degraded (pre-recolonisation and pre-restoration) conditions. This yielded 25 initial simulations (5 food webs x 5 landscapes), 25 recolonisation simulations without restoration and 750 restoration simulations (25 initial simulations x 6 restoration sequences (5 random + 1 clustered) x 5 restoration levels (1 to 5 patches restored)). This amounted to a total of 800 simulations in which 15 local food webs (number of patches in the landscape) were modelled, for a total of 12,000 food webs.

All simulations were performed on python version 2.7.5 available at https://www.python.org/ using modules *numpy* version 1.26.4 (Harris *et al*., 2020), *scipy* version 1.12.0 (Virtanen *et al*., 2020) and *pandas* version 1.5.1 (The pandas development team, 2024) (see the *Code availability* section for code and instructions to replicate the simulation experiments).

### Analysis of experimental outcomes

To assess the outcome of restoration in our model simulations, we recorded species’ biomasses, species’ traits, and local (patch-level) food web properties before and after recolonisation and/or restoration. All metrics were extracted from the final communities that had reached “stable” population dynamics (see *Initial population dynamics* section for stability criteria).

#### Number of recolonisers

The number of successful recolonisers was evaluated against the species composition in the initial local community before recolonisation was allowed. A recoloniser was considered successful if it was absent from the initial community and its biomass was larger than the extinction threshold (1e^-8^ g) after population dynamics had stabilised.

We categorised recolonising species into four trophic levels: plants (basal species with no resources), herbivores (species feeding exclusively on plants), intermediate species (feeding on a combination of plants and herbivores), and top consumers (species with no predators). In addition, we measured recoloniser generality (number of resource items consumed) and vulnerability (number of predators). All feeding traits were recorded from the regional 100 species food web.

#### Species persistence

We recorded local species persistence as the proportion of extant species (whose biomass was larger than the extinction threshold) among the potential maximum number of species from the regional pool (100 species). Persistence was also recorded per trophic level as the proportion of extant species among the number of potential species from this trophic level in the regional pool.

#### Food web structure

We recorded local community features, including food web metrics, before and after restoration. These included (1) species richness, (2-5) number of species per trophic level (basal, herbivore, intermediate, top), (6) standard deviation of generality (number of feeding links) and (7) vulnerability (number of predatory links) of the food web, (8) total number of links per species (feeding + predatory links), (9) mean food chain length, calculated as mean number of edges between all pairs of resource and top consumer species in the food web, and (10) network modularity (Eq. 11). Modularity is a network metric that quantifies the extent to which nodes within the same group of nodes (i.e. module) are more densely connected to each other than to nodes in other modules. Modularity takes positive values when there are more edges between species within modules than we would expect by chance (Newman, 2010). We computed modularity following Clauset *et al*. (2004) (Eq. 11) using the *networkx* module in *python*.

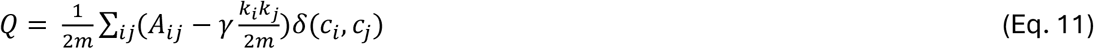

Where m is the number of links in the network, A is the adjacency matrix of interactions between nodes, *k*_*i*_ and *k*_*j*_ are the degrees of species i and j, *γ* is a resolution parameter (kept to 1 here), *δ*(*c*_*i*_, *c*_*j*_) is 1 if i and j are in the same module and 0 otherwise. Module partitions were obtained using the Clauset-Newman-Moore greedy modularity maximization algorithm (Clauset *et al*., 2004) and then finding the partition that maximise modularity, starting with each node being in its own module and joining pairs of modules until no more increase in modularity is achieved.

#### Comparing clustered and scattered restoration scenarios

We quantified how restoration affected food web structure by comparing the distribution of food web metrics before and after restoration/recolonisation and at different restoration levels (number of patches improved).

To compare the outcomes of clustered vs scattered patch restoration scenarios, we plotted mean biomass, species richness and the different facets of food web structure defined above against number of improved patches for each scenario with 95% confidence intervals across landscapes and food web replicates. As for the effect of restoration, we quantified the differences between restoration scenarios by comparing the distribution of food web metrics in clustered and scattered landscape at each stage of restoration from the initial stage to 5 patches restored.In addition, we computed the Shannon alpha diversity of each patch 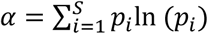, where *p*_*i*_ corresponds to the proportional abundance of each species 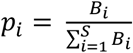.

We also explored differences between clustered and scattered scenarios in terms of species accumulation curves. Patch isolation in meta food webs promotes beta diversity (Ryser *et al*., 2019) and thus, we expected that the central and more connected patches that are improved in the clustered scenario would be more similar in terms of species composition, hence benefiting fewer species than the scattered scenario. We plotted species accumulation curves for patches to be restored in initial simulations, respecting the sequence of restoration in the clustered and 5 scattered scenarios and then retrieving the same curves for restored patches after restoration. In this way, we were able to explore how restoration of clustered or scattered habitat patches modulates the outcome of restoration. For example, we expect that species accumulation curves would plateau faster for clustered than for scattered scenarios.

#### Evaluating the relative importance of initial conditions and landscape properties

We wanted to understand whether initial food web and landscape properties could predict the number of successful recolonisations across trophic levels. We ran random forests to predict the number of recolonisers across trophic levels from 10 food web metrics of the initial local communities and 5 landscape or patch characteristics. Landscape characteristics used included number of improved patches, Euclidean distance to closest improved patch, Euclidean distance to source (top-left) patch, summed Euclidean distance to other patches and restoration type (scattered or clustered). From the 10 food web metrics described above we kept 8 that seemed most relevant for predicting recolonisations: species richness, connectance, standard deviation of generality and vulnerability, modularity, and number of top, intermediate, herbivores and basal species.

Random forests are machine learning methods used to predict categorical outcomes from complex relationships among many variables by creating multiple decision trees based on bootstrapped samples of the input data. For example, if we wanted to predict the number of recolonisations based on the distance from the source patch: from the root of the tree, the first question is “How far is the source patch?”. If the source patch is less than e.g. 0.2 units away, there will likely be more recolonisations. And from there the same process is followed for further predictor variables. We trained random forests on 80% of the data and prediction accuracy was tested on the remaining 20% left out. We set the number of trees in the forest to 200 with a maximum tree depth of 10 to prevent overfitting and used a random seed to ensure replicability. From the random forest outputs, we extracted prediction accuracy (number of correct predictions/total number of predictions) and variable importance for all predictors. Variable importance is a measure of accuracy loss when each variable’s values are randomly permutated (shuffled). To report the sign of the relationship, we extracted the coefficient estimate from a generalised linear model with a poisson distribution of the relationship between number of recolonisers and the interaction term between trophic level and drivers of recolonisation (listed above). Random forests were run with the python package *sklearn* (Pedregosa *et al*., 2011).

## Results

Before recolonisation or restoration, our 375 patches (25 simulations x 15 patches) had on average 35.6 (sd = 8.5) species with an average of 3.4 trophic links (sd = 0.9). Mean food chain length was on average 3.6 (sd = 0.8). Enabling recolonisation in these landscapes without any restoration caused a slight increase in species richness to an average of 36.0 (sd = 8.5).

### Habitat restoration increases biodiversity and biomass across trophic levels

Patch improvement, implemented by increasing the growth rate of basal (e.g. plant) species three-fold in improved patches, effectively increased the amount of energy travelling to higher trophic levels. This allowed increases in mean biomass and in the number of species per patch across the whole landscape from an average of 35.6 (sd = 8.5), up to 39.7 (sd = 10.0) species when 5 patches were improved (Fig. S1.1). Thus, on average, improving 1/3 of the landscape allowed for the recolonisation of 4.1 additional species per patch compared to the landscape before recolonisation and restoration.

Recolonising species were found mainly at intermediate (on average 3.25 new intermediate species after all 5 patches were improved) and, to a lesser extent, top positions (on average new 0.48 species) in the food web (Figure 2B). The total biomass of intermediate species increased by 45.45% while the total biomass of top species more than tripled, increasing by 255% after 5-patch improvements (Figure 2C). Thus, despite gaining fewer species, the mean biomass of top species increased sharply with patch improvement, indicating that top species became more abundant whilst intermediate species became more numerous (Figure 2B, C, E, F). In addition, total and mean biomass of top species increased slowly with the first patch improvement and then picked up and increased linearly with subsequent improvements (Figure 2C, D).

**Figure 2:**
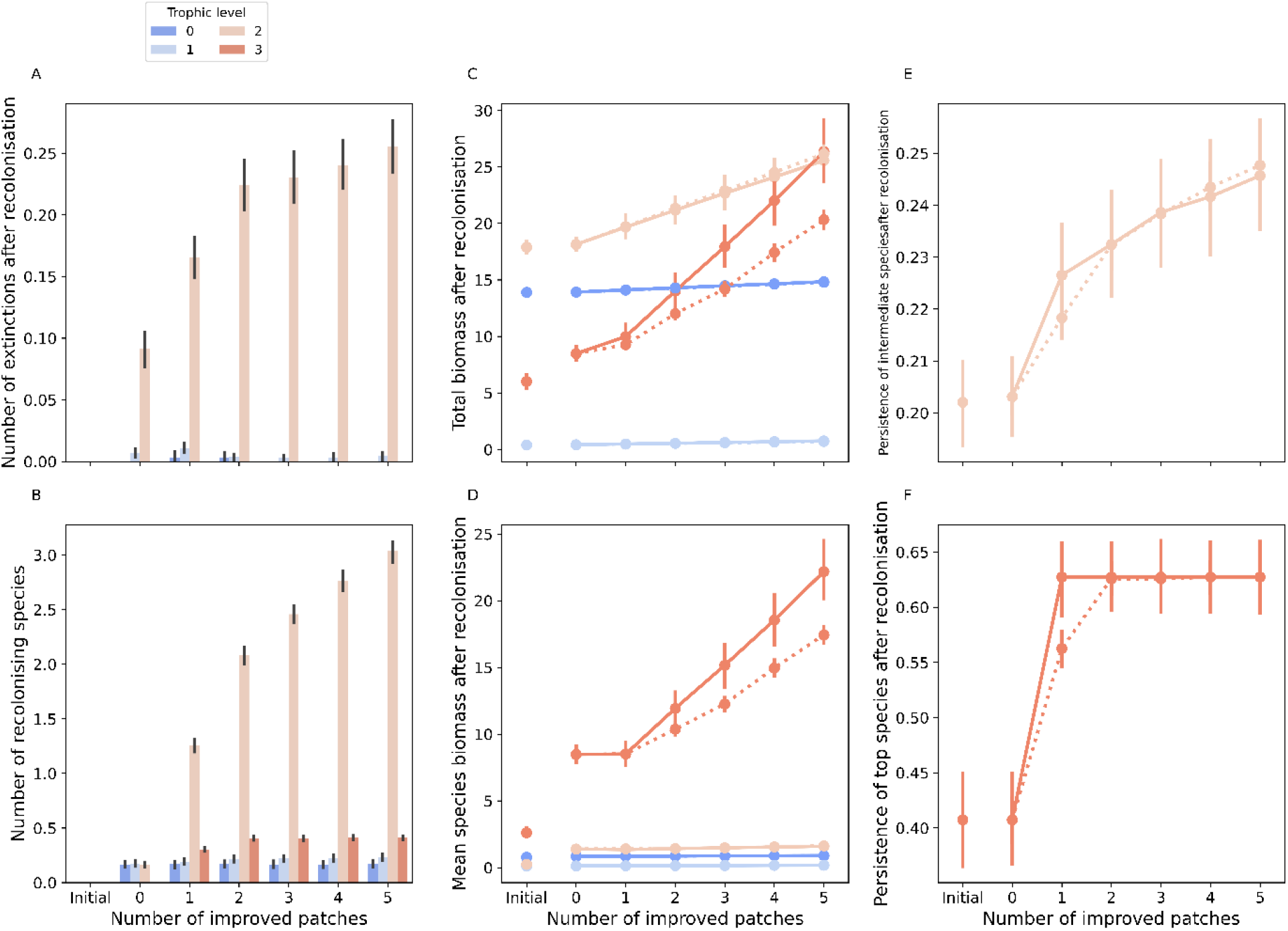
Patch improvement increases local biomass, species persistence and number of intermediate and top species in meta food webs. A and B show the mean number of local extinctions and recolonisations across trophic levels under both clustered and scattered scenarios. C and D show the increase in total biomass and mean species’ biomass of species separated into four trophic levels. E and F show mean persistence of intermediate and top species (persistence of plants and herbivores remained rather constant and high – see main text). In panels A-B bars represent mean number of extinctions and recolonisation per trophic level, while error bars represent 95% confidence intervals around the mean. In panels C-F: points represent the average value of the given variable, and error bars the 95% confidence interval around it. Solid lines represent results from the clustered scenario where the five improved patches were clustered in the middle of the landscape. Dotted lines correspond to results from the scattered scenario where improved patches were randomly placed in space. Across all panels, colours represent trophic levels from 0 to 3, respectively plants (0 - dark blue), herbivore (1 - light blue), intermediate species (2 - light red) and top species (3 - dark red). Initial results are for “degraded” simulations before recolonisation. 0 patch improvement corresponds to the initial simulations after allowing for recolonisation.

Importantly, we found that top species were only able to recolonise the landscape after at least one patch was improved (Figure 2B,F). Persistence of herbivores and plants before recolonisation was already very high (herbivores: mean = 79.0%, sd = 24.6%; plants: mean = 90.4%, sd = 0.8%) and thus very few recolonisation events were recorded on average for those species, therefore the average number of recolonisation events from these species stayed constant (Figure 2B). We observed rather few extinctions throughout all “non-initial” simulations (mean number of extinctions = 0.02, sd = 0.20), and mainly in herbivores (mean = 0.01, sd = 0.09) and intermediate species (mean = 0.08, sd = 0.34) (Figure 2A). Lastly, the increase in persistence of intermediate and top species with restoration was not linear, increasing faster with the improvement of the first patch and subsequently slowing down (Figure 2E), and even plateauing for top species (Figure 2F).

Beyond species richness, patch-improvement caused changes to the topology of local food webs. Overall, food webs became more complex (both in species and interactions; Fig. 3A-B) with patch improvement, resulting in longer food chains (Fig. 3D). Species became more generalists on average, with more links per species (Figure 3E). Increases in mean body mass (Fig. 3C) were largely driven by the successful recolonisation of top predators. When accounting for diversity in species richness as well as biomass (Shannon diversity), biomass became more evenly distributed across species within communities along the gradient of patch improvements (Figure 3F).

**Figure 3:**
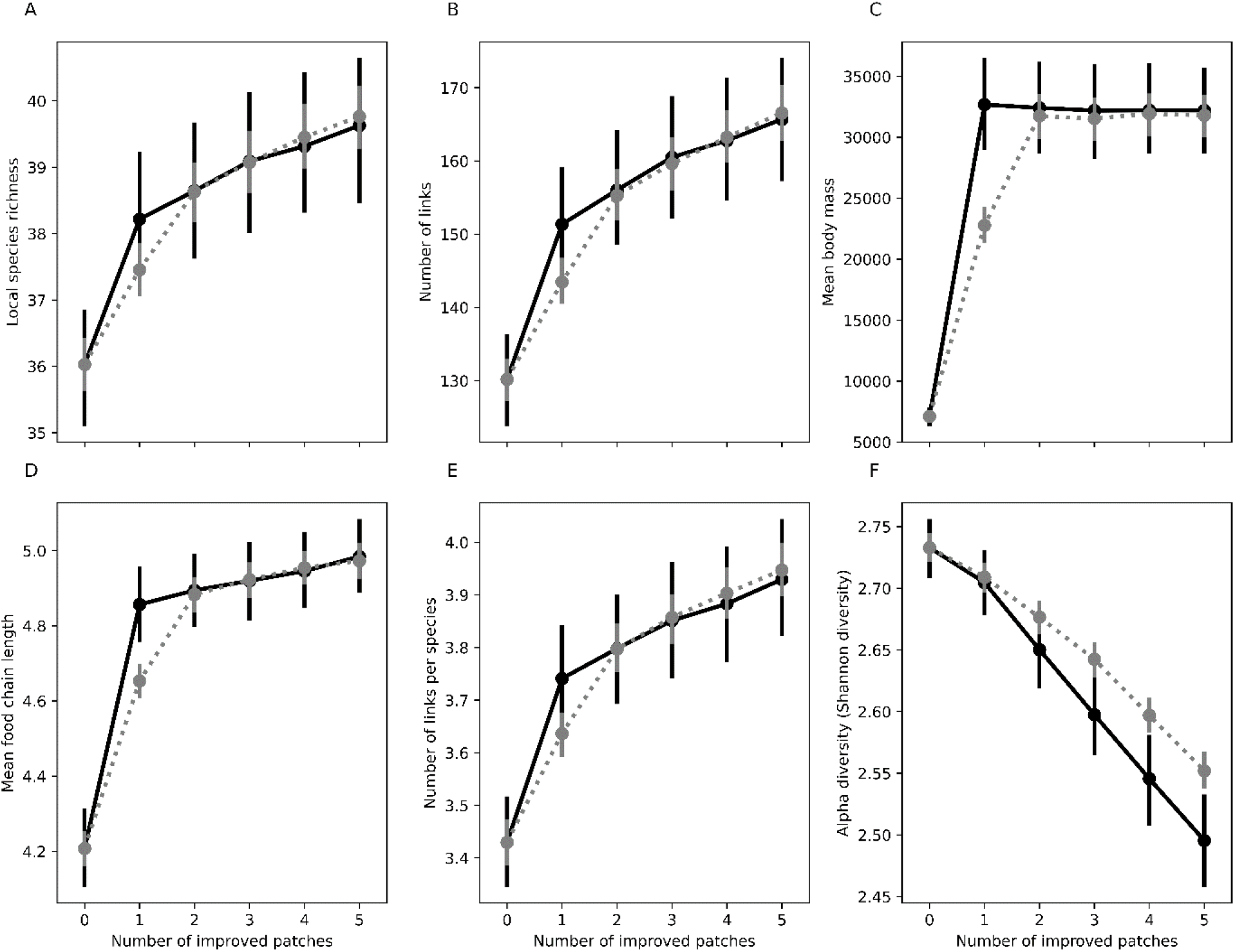
Patch restoration improves local food web structure and evens species composition. We show how the average patch-level food web structure or community feature (points) and 95% confidence interval (error bars) vary with number of improved patches. Results are for all patches (restored and non-restored). Solid lines represent results from the clustered scenario where the five improved patches were clustered in the middle of the landscape. Dotted lines correspond to results from the five different scattered scenarios where improved patches were randomly scattered in space. 0 patch improvement corresponds to the initial simulations after allowing for recolonisation. A larger number of improved patches resulted in an increase in food web complexity and mean food chain length. The differences between the clustered and scattered scenario is mainly apparent when only one patch is improved.

### Recolonisation is driven by landscape and food web features

To unveil the mechanisms behind differences in number of successful recolonisations across patches, and trophic levels, we ranked the importance of pre-restoration conditions of the meta food webs in predicting the number of successful recolonisers using random forests classification trees. We found that the most important predictors of recolonisation within plants, herbivores and intermediate species were spatial attributes of the patches across the landscape (Fig. 4). For plants and herbivores, the single most important characteristic modulating the number of recolonisers was distance to the source patch (plants: e = −32.29, p < 0.001, herbivores: e = −10.44, p < 0.001) (see Table S1 and Figs. S1.2 and S1.3 for other drivers of recolonisation). Indeed, as we see in the inset graphs of Fig. 4A and B, successful colonisers were predominantly found in patches close to the invaded one, while patches further away generally experienced no colonisation events. Finally, herbivores recolonised more readily landscapes where the modularity of the food web was low (e = −4.65, p < 0.05) but also those food webs with less herbivore species (e = −0.52, p < 0.001), but more species in general (e = 0.49, p > 0.05), higher connectance (e = 61.43, p < 0.001) and with larger variation in diet breath (e = 31.91, p < 0.001). The spatial attributes of patches together with local food web structure allowed the prediction of the number of plant and herbivore recolonisations with an accuracy of 99% and 97%, respectively.

**Figure 4:**
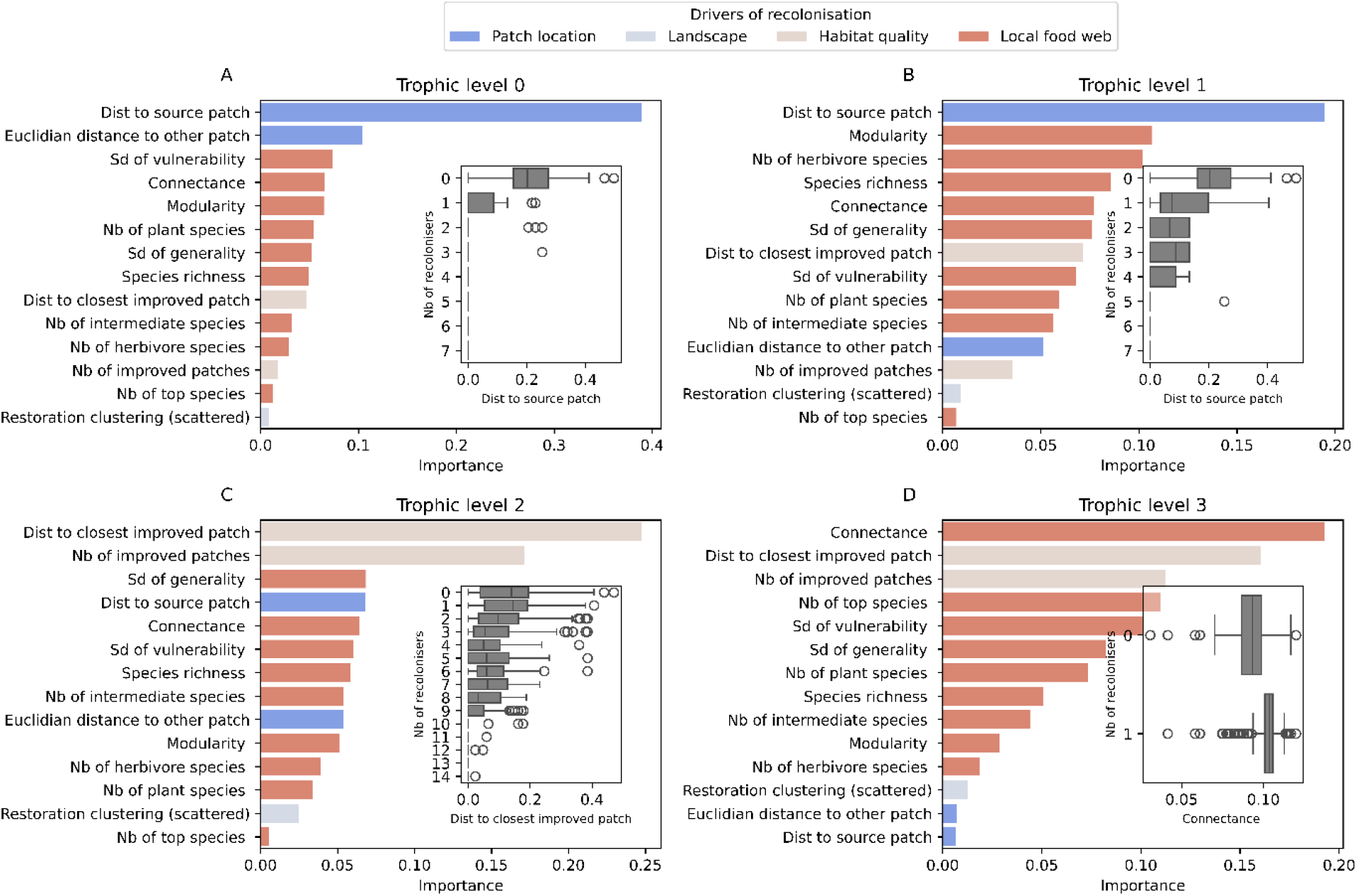
Pre-restoration conditions of the food web and the landscape of patches drive the number of recolonisations, but drivers differ across trophic levels. Plots show variable importance from random forest analysis of initial patch and food web characteristics in predicting the number of local recolonisations per trophic level in simulations from 0 to 5 patch restorations. Bars represent variable importance (evaluated from the loss of predictive accuracy resulting from permutating each predictor variable – see Methods) in predicting the number of successful recolonisation events across trophic levels. Initial conditions (before invasion and restoration) were measured at the local (i.e. patch) level and include (1) patch location context metrics (dark blue bars) such as distance to invaded patch or connectivity of the patch, (2) landscape attributes (light blue bars): whether restoration was clustered or not, (3) habitat quality related information (orange bars) such as distance to improved patch and number of improved patches and (4) local food web structural properties (orange bars). Inset boxplots represent median, 25% and 75% quantiles of the highest ranked predictor variable across different numbers of recolonisations. They illustrate the relationship between number of recolonising species and the most important predictor variable for that trophic level (ranked by variable importance in random forests). All other relationships are illustrated in Figs. S1.2-5.

Recolonisation of the landscape by intermediate and top species on the other hand, was best described by landscape features associated with patch quality. Thus, proximity to an improved patch (e = −3.41, p < 0.001) as well as the number of improved patches (e = 0.13, p < 0.001) across the landscape were the main predictors of recolonisation by intermediate species (Fig. S1.4), albeit yielding a lower accuracy of 64%. Top species recolonisation was best predicted by a combination of food web and landscape characteristics related to habitat quality such as distance to, and number of improved patches. Indeed, as observed in Figure 2F, top species were only able to recolonise the landscape after at least one patch was improved (see also Fig. S1.5).

Random forest classification accuracy was 94% for top species. The most important food web property in determining recolonisation success for top species (Fig. 4D) was local food web connectance, with increased recolonisations on less connected food webs (e = 20.56, p > 0.05). The number of top species (e = −1.89, p <0.001), the standard deviation of vulnerability (e = −15.71, p <0.05), and the standard deviation of generality (e = 66.18, p < 0.001) were also found to be important drivers of recolonisation for top species, with many recolonisations observed on food webs with larger variability in diet breadth, less variability in number of predators and in patches with fewer top species (see Fig. S1.5 to visualise effect of all drivers on recolonisation of top species).

### Spatial patch configuration modulates the outcome of restoration across clustering scenarios

Gains in species richness when only 1 patch was improved were on average smaller in the scattered scenario, where the improved patch could be located anywhere in the landscape, than in the clustered scenario, where the patch was central to the landscape (Figure 2E,F and Fig. S1.6). These smaller species gains for the scattered scenario for the single improved-patch case were linked to the location of the improved patch. A more connected patch (closer to other patches) gained more top and intermediate species on average compared to a more isolated patch (Fig. S1.6).

This trend, however, disappeared beyond 2 patches improved (Figure 2E,F). Species area curves indicate that the number of species benefitting from restoration in the scattered scenario slightly exceeded, on average, the number of species benefitting from restoration in the clustered scenario beyond 2 patches improved (Fig. S1.7). Nonetheless, beyond 2 patches improved differences were small between scenarios. Differences between the scattered and clustered scenarios in terms of food web metrics mirror those of species richness, where the location of the restored patch in the 1-patch case influenced restoration outcome, but the difference vanished when more patches were improved (Figure 3).

On the other hand, biomass increase with patch improvement in the scattered scenario was significantly slower than in the clustered scenario beyond one patch improved. The absence of difference in biomass between the two scenarios in the 1-patch improvement but subsequent increase, suggests that clustering of restored patches has potential beneficial effects on the ecosystem beyond increases in species richness or network metrics, with the observed additive (linear) effect on biomass increase (Figure 2C and D) revealing increases in ecosystem productivity.

## Discussion

“The practice of ecological restoration requires a high degree of ecological knowledge that can be drawn from practitioner experience, Traditional Ecological Knowledge, Local Ecological Knowledge […], and scientific discovery.” (Gann *et al*., 2019). Although restoration ecology is given more and more attention from the scientific community, it still lacks important links with ecological theory (Zanden *et al*., 2016). In this study, we aimed to contribute to this knowledge base through a realistic multi-layer modelling framework aimed at informing ecological restoration by coupling trophic interactions and dispersal dynamics. These two elements are key drivers of community assembly and considering them in synergy enabled better understanding of the restoration process.

We found that restoring habitat-patches in a landscape improved community composition and food web structure by enabling the recolonisation of higher trophic-level species. Interestingly, recolonisation from species from the regional food web did not prompt major extinction events, in contrast to what has been found in theoretical studies which predict “large biodiversity loss” in complex communities from invasions (e.g. Lurgi *et al*., 2014) as well as empirical meta-analyses reporting 21% species richness decline in aquatic and a 27% decline in terrestrial habitats due to invasions (Mollot *et al*., 2017). Our study differed from previous studies focusing on invasions (e.g. Romanuk *et al*., 2009; Lurgi *et al*., 2014; Häussler *et al*., 2021; Sentis *et al*., 2021) in that our recolonisers were not exotic species nor particularly strong competitors with extreme traits. Rather, they were species that were not able to survive under the initial poorer habitat quality conditions for lack of enough resources to maintain viable populations.

By providing influxes of individuals from restored patches, source-sink dynamics (Brown & Kodric-Brown, 1977) through rescue effects enabled the persistence of otherwise absent species in surrounding non-restored patches. This mechanism is responsible for the landscape-wide improvements of both species richness and food web metrics (mean food chain length, connectivity) we observed from restoring just a portion of the landscape. Overall, restoration benefitted not only food web structure in restored patches but also surrounding patches. In general, habitat heterogeneity in meta food webs enables higher overall biodiversity compared to their homogeneous counterparts (Ryser *et al*., 2021). These rescue-effects have been observed in natural systems where influxes of colonisers from non-disturbed habitats were found to dampen the effects of a disturbance in a connected patch in a two-habitat experimental systems (Altermatt *et al*., 2011).

### Dispersal limitation varies across trophic levels

Local conditions of disturbed habitats are known to influence the order of recolonisation in perturbed landscapes through priority effects, where initial colonisers modulate resource availability thereby facilitating or hindering the establishment of later successional species (Weidlich *et al*., 2021). To explore whether initial community characteristics modulated recolonisation potential, we assessed the relative importance of improvement levels, patch location and initial food web structure in predicting recolonisation of restored and non-restored landscape across different trophic levels. We found that patch-location and landscape improvement levels were better predictors of the number of successful recolonisation events across all trophic levels except top species than food web structure. Similarly to (Romanuk *et al*., 2009; Smith-Ramesh *et al*., 2017), who found different roles of initial conditions depending on the trophic level being invaded, we found that different trophic guilds were sensitive to different meta food web features. Further, in line with (Häussler *et al*., 2021) who found that invasion was modulated by an interplay between invaders’ dispersal ranges and nutrient supply, we found that differences in recolonisation between trophic levels were underpinned by the relative energy requirements versus dispersal ability of the recolonising species. Thus, plants and herbivores, in general poorer dispersers in our model, were mostly limited in their recolonisation ability by the distance to a source population, meanwhile, restoration levels and habitat quality only played a secondary role for lower trophic levels. This agrees with the empirical findings of e.g. Winking *et al*. (2014) and Kitto *et al*. (2015) who looked at the recovery of invertebrate communities in mine-impacted waterways in New Zealand and in old sewage channels in Germany, respectively. Both studies found that proximity to a source stream was an important driver of community composition. Havel *et al*. (2002) studied the spread of an invasive species of zooplankton in Missouri (USA) lakes over seven years and found similar results. Interestingly, they also found that lake productivity did not affect invasion success and thus hypothesised that other forms of biotic control besides energy limitation, might be responsible for modulating invasion success. Empirical evidence thus confirms the importance of proximity to source patch for those lower trophic levels.

On the other hand, recolonisation by intermediate and top species who have better dispersal capabilities but high energy requirements, were rather driven by distance to restored patch and restoration levels, as well as food web structure, rather than dispersal limitation. Although, it is likely that all species are limited in their dispersal range either by their dispersal ability or by behavioural traits, they might be so to different spatial extents (hundreds of meters for bees (Ricketts, 2004), dozens of kilometres for seabirds (Buxton *et al*., 2014)). Thus, for example, even in seabirds, supposedly very good dispersers, one of the major drivers of recolonisation success after the removal of invasive mammals from New Zealand islands is the proximity to a source population (Buxton *et al*., 2014).

This highlights the different scales at which species operate, and the importance of considering species’ dispersal habits at all trophic levels in relation with habitat connectivity and notably, distance to source populations in restoration projects.

### Mechanisms behind the increase in number of intermediate species and biomass of top predators

Pimm’s productivity hypothesis states that food webs should increase in length with increasing productivity (Pimm, 1982). The amount of energy entering the food web constitutes a limiting resource in the number of trophic levels and species an area can sustain (Pimm & Kitching, 1987). Because in our system, the first two trophic levels are almost “saturated” i.e. have high persistence even before restoration, the main food web improvements observed are in subsequent trophic levels of intermediate and top species. As predicted by the energy limitation hypothesis (Lindeman, 1942; Elton, 1958), we found that the initial amount of resource or energy available in our system limited the number of species and trophic levels it could sustain. Increasing nutrient input in selected patches enabled the recolonisation of intermediate and top species from the regional pool, improving species persistence, a key metric that conservation ecology usually aims to maximise (Török & Helm, 2017). Notably, no top species were able to recolonise the landscape before restoration.

Improving increasing portions of the landscape caused linear increase in the total biomass of intermediate and top trophic levels and translated into more numerous intermediate species. However, number of top species increased a lot slower, only gaining species incrementally. Rather it was top species’ mean biomass that increased with patch improvement. Indeed, the amount of energy necessary to sustain top species is much higher than that of smaller species given their higher body size, metabolic rates and overall energy requirements (Arim *et al*., 2007). In line with our expectations, this step-like increase in the number of top species suggests that while biomass increases linearly with energy input, quite a high biomass increment in prey items from below trophic levels is necessary to sustain new top species in the system. Thus, the recolonisation of a higher trophic level species occurred only if sufficient biomass had been gathered in the trophic levels below to sustain this species. It follows that the distance to, and number of, improved patches would underpin recolonisation of top and intermediate with higher energy requirements. Thus, in natural ecosystems we should expect patches closer to improved areas to benefit from the spillover of species from the restored patch, themselves benefiting from increased nutrient input, causing higher emigration fluxes.

Although recolonisation by top predators was relatively rare in our simulations, our results suggest that further patch improvement enabled more specialist top and intermediate species to colonise (Fig. S1.8). This highlights the importance of quantitative approaches to restoration aimed at preserving rare species that might be limited by their degree of specialisation and the availability of specific prey. These methods could help forecast “how much” restoration is necessary to sustain those rare species. Although in empirical systems, the major threat to predator reintroduction is anthropogenic disturbances, e.g. illegal hunting of Lynx populations in Central Europe (Heurich *et al*., 2018) or bear mortality in North America (Gantchoff *et al*., 2020), and reducing these threats are probably the first step to encourage the recolonisation of key top predators. Our results show that conditions inherent to the ecosystem, notably aspects of productivity and connectivity, have the potential to encourage the return of those species provided other aspects of anthropogenic disturbance are also mitigated.

In addition to patch location and improvement levels, initial food web structure was an important predictor of recolonisation by top species, and to a lesser extent herbivore recolonisation. Herbivores tended to recolonise communities with less herbivore species. Meanwhile, similar to the findings of Romanuk *et al*. (2009) for carnivore invaders, top species mostly recolonised food webs with fewer top species. Overall, available niche space seemed to be an important criterion for invasion success for herbivores and top species.

The expectation is that high connectance should confer biotic resistance and reduce available niche space for invaders e.g. (Romanuk *et al*., 2009; Galiana *et al*., 2014; Sentis *et al*., 2021). After accounting for collinearity with other food web metrics (e.g. species richness) we indeed found that more top species recolonisation occurred in landscapes with smaller food web connectance. However, as reviewed by (Frost *et al*., 2019), in the invasion literature the relationship between connectance and invadibility remains unclear and they suggest that this relationship should be considered in relation to species diversity and trophic level of the introduced species. Our food webs followed the constant connectance hypothesis, thus connectance varied independently from species richness in our system. We also found that less speciose food webs favoured top species recolonisation (Fig. S1.5), in agreement with the biotic resistance hypothesis that posits that species-rich communities are more resistant to invasion (Elton, 1958).

Beyond mechanistic effects of food web structure, the recolonisation of seabirds in New Zealand was found to be favoured by bird species richness in recolonised environments (Buxton *et al*., 2014). It was proposed that islands with more ecological similar species might encourage seabirds to establish there. Although those behavioural aspects were not considered in this study, they could also play a role in modulating recolonisation success in real-world landscapes.

### Restoration of well-connected patches is a good policy when only a small portion of the landscape can be restored

The improvement of just one patch had disproportionate positive effects on species richness and food web structure compared to subsequent improvements (i.e. more than one patch). We hypothesise that this observation could be linked to the species area relationship (SAR). Beta diversity in our landscapes was relatively low, thus, one single patch improvement enhanced a large fraction of the regional species pool. Subsequent improvements only captured a handful more species, thus having smaller relative impact on the overall community.

In addition, the location of the first restored patch strongly influenced the outcome of restoration. To understand this difference in improvement outcome in the first instance of patch improvement, we explored the role of patch location and connectivity. Landscape connectivity has been found to enhance animal-dispersed plant diversity in areas surrounding connected patches compared to unconnected patches in corridor experiments (Brudvig *et al*., 2009). Similar spillover effects benefiting fisheries around marine reserves have been abundantly documented (Gell & Roberts, 2003) as well as patterns of increased pollinator activity in ecosystems in proximity to tropical forest fragments (Ricketts, 2004). We found that restoring a better-connected patch in the first instance enabled the recolonisation of 1 additional intermediate species compared to that of an isolated patch. This is a result of spillover and rescue effects (Brown & Kodric-Brown, 1977; Gravel *et al*., 2010; Häussler *et al*., 2021; Ryser *et al*., 2021) from restored patch to neighbouring ones. Thus, if only a small portion of the landscape can be restored, strategically restoring well connected patches should improve restoration outcomes through spillover effects to surrounding patches.

### Fragmentation modulated biomass change but not species richness

The SLOSS (Single Large Or Several Small) problem is a key discussion around reserve planning, asking whether for the same total area, a single large (SL) reserve would encompass more species richness than several smaller (SS) reserves. Early studies concluded that SL reserves should be favoured over SS (Diamond, 1975) based on Island Biogeography theory (MacArthur & Wilson, 1967), and this was adopted as a gold standard for conservation efforts (Fahrig, 2020). Nowadays, empirical studies usually address the SLOSS problem by comparing species accumulation curves (SACs) for reserves ordered from the smallest to the largest versus the largest to the smallest, and comparing curves at equal total area (Quinn & Harrison, 1988). If the smallest to largest curve is consistently above the largest to smallest curve, then studies remain in favour of SS over SL, while if curves cross results are inconclusive (Fahrig, 2020). A review by (Fahrig, 2020) showed that the literature on the SLOSS dichotomy mostly agrees that several small (SS) reserves should maximise species richness over single large (SL) ones.

Here, we find inconclusive evidence in favour of either SL or SS. Despite heterogeneous species composition in the initial landscape and evidence that a scattered restoration sequence could potentially capture more species, the difference between species accumulation curves was not significant across restoration experiments. Our results suggest that provided that the initial habitat heterogeneity is quite low, and thus species composition is rather homogeneous across the landscape, improving patches in a clustered or scattered manner makes little difference provided a larger enough portion of the landscape is improved (here larger than 13% - 2 patches out of 15 patches). We hypothesise that, had the initial landscape been more heterogeneous or larger with higher beta diversity, the scattered scenario (SS) would have shown improved outcomes compared to the clustered distribution (SL) in terms of species diversity and food web structure by leveraging on source-sink dynamics across patches (Lasky & Keitt, 2013). Further research inspired by our work should focus on obtaining a better understanding of landscape heterogeneity on restoration.

Despite species richness not significantly differing between clustered and scatter scenarios, the biomass increase of top species did show marked differences. Clustered restoration scenarios maximised the regional biomass increase of top species, because their initial biomass was significantly higher in centre patches than in periphery patches. In the initial (i.e. pre-restoration) landscape, middle patches (those improved in the clustered scenario) held on average larger biomass than peripheral patches just by virtue of spatial dynamics favouring short distance dispersal at that scale. Improving those patches thus disproportionately increased biomass of top species. Biomass increased linearly, suggesting that increasing habitat quality across the landscape has an additive rather than multiplicative effect on species biomass.

Increasing biomass but not species diversity could have implications for community stability. Diversity is usually considered a stabilising feature for communities. As Yachi and Loreau (1999) note: “according to the insurance hypothesis, biodiversity insures ecosystems against declines in their functioning because many species provide greater guarantees that some will maintain functioning even if others fail”. However, scenarios in which increasing nutrient input beyond a critical threshold leads to increases in top species biomass but not number of top species can lead to the paradox of enrichment (Rosenzweig, 1971), where strong top-down control results in biomass fluctuations, leading to destabilising fluctuations and loss of diversity. This has been shown in meta food web models like the one used here, where beyond a certain nutrient input, species diversity decreased despite dispersal and drainage effect (Ryser *et al*., 2021).

### Limitations and perspectives

Empirical work has shown that manipulating the order of arrival of plant species can alter the outcome of restoration (Popp *et al*., 2017). This order can determine plant dominance, affect species’ biomass and ultimately influence overall patch composition. In our study, we allowed all absent species to recolonise at once, meaning that they all competed for a limited set of resources, and only the species that were best suited for recolonisation given the resources available were able to establish. Varying the order of recolonisation or allowing only certain species to recolonise may have altered the outcome of restoration. Future research could explore how varying this order of recolonisation affects community assembly after restoration, and whether this could be used to optimise species persistence.

In addition, it would be interesting to further explore the role of the species-area and network-area relationships (Galiana *et al*., 2018) on restoration, and whether they could be effectively used to develop restoration strategies aimed at improving food web structure. Notably, it is possible that in a landscape with higher heterogeneity the scattered scenario of improvement would exceed that of the clustered scenario beyond a few patch improvements. This would effectively inflate the differences between scattered and clustered scenarios in our simulations.

Our exploration does not explicitly vary patch size, rather, it varies the number of high-quality patches and their clustering. However, it would be interesting to explore patch-size explicitly, given that patch size is at the root of the Theory of Island Biogeography (TIB) and the SLOSS debate. The TIB posits that large patches should have lower extinction rates and higher immigration rates, and thus, patch size is often found to be an important driver mean food chain length in empirical systems (Doi *et al*., 2009; Takimoto & Post, 2013). For example, (Ryser *et al*., 2024) explored the Island Species Area Relationship in the context of meta food web models, explicitly modelling patch size, population density and varying dispersal fluxes between different sized patches. Similar approaches could be used to explore the interaction between patch size and habitat heterogeneity in a restoration context.

### Concluding remarks

Conservation goals are usually set in terms of proportion of land set aside for protection. Aichi Target 11 from the Convention on Biological Diversity (CBD) states that “by 2020, at least 17 per cent of terrestrial and inland water, and 10 per cent of coastal and marine areas […]” should be conserved. Our results suggest that indeed, restoring larger portions of the landscape yields systematically better results than restoring smaller parts of the landscape, but also that if restoration of only a small portion of the landscape is possible - in agreement with previous literature (Brudvig *et al*., 2009) - restoring more connected patches could optimise restoration outcomes. This will in turn yield disproportionately positive results in habitats well-connected to the surrounding landscape. In addition to increasing the number of patches restored, we found that patch location was an important modulator of restoration success across trophic levels in meta food web models. Thus, integrating considerations about the dispersal ability and resource requirements of the species that we wish to bring back in restoration projects should constitute a priority if we wish to promote the recolonisation of those rare species. Furthermore, in addition to strategic planning of the location of restored/protected patch, our findings support landscape restoration as an effective way to bring back and sustain top predators and more specialist top predators, although this remains to be further tested. Finally, our results suggest that, in a sufficiently homogeneous and well-connected landscape, single large or several small reserves should yield equivalent results in terms of species richness but also food web metrics beyond rough threshold of ~13% landscape restoration.

## Code availability

Code and instructions to replicate this analysis can be found: https://github.com/LucieTp/Patch-restoration-Meta-food-webs

