## Supplementary Figures for "Habitat restoration promotes recolonisation by extirpated species in model meta food webs"

Supplementary material for the paper: ***Habitat restoration promotes recolonisation by extirpated species in model meta food webs***

**Authors:** Lucie Thompson^1,*^, Miguel Lurgi^2,*^

**Affiliations:**

^1^ Centre d'Ecologie et des Sciences de la Conservation (CESCO), Muséum national d'Histoire naturelle, Centre National de la Recherche Scientifique, Sorbonne Université, CP 135, 57 rue Cuvier 75005 Paris, France

^2^ Department of Biosciences, Swansea University, Singleton Park, SA2 8PP, UK.

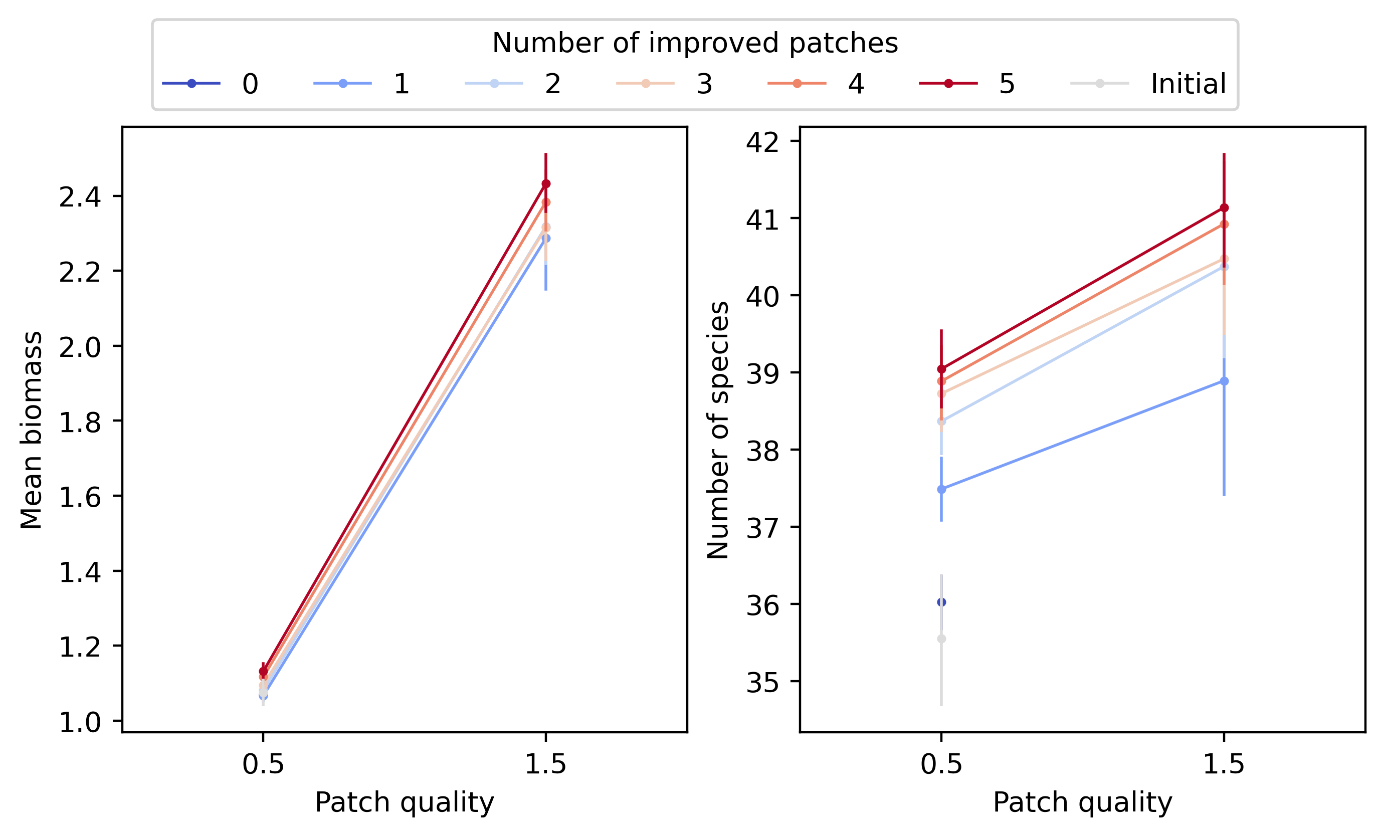

***Figure S1.1: Difference in biomass and species richness between restored (1.5) and low quality (0.5) patches coloured by number of improved patches.*** Points are means across replicate food webs, landscapes and clustering scenarios. Error bars show the 95% confidence intervals around the mean. Plots illustrate the effects of restoration on mean species biomass (after stabilisation of population dynamics) and number of species per patch. Lines linking points between the low patch quality (0.5) and high patch quality (1.5) were added for legibility. Initial simulations (grey points) are for landscape before recolonisation and restoration. 0 patch improvement is for simulations before restoration but allowing recolonisation.

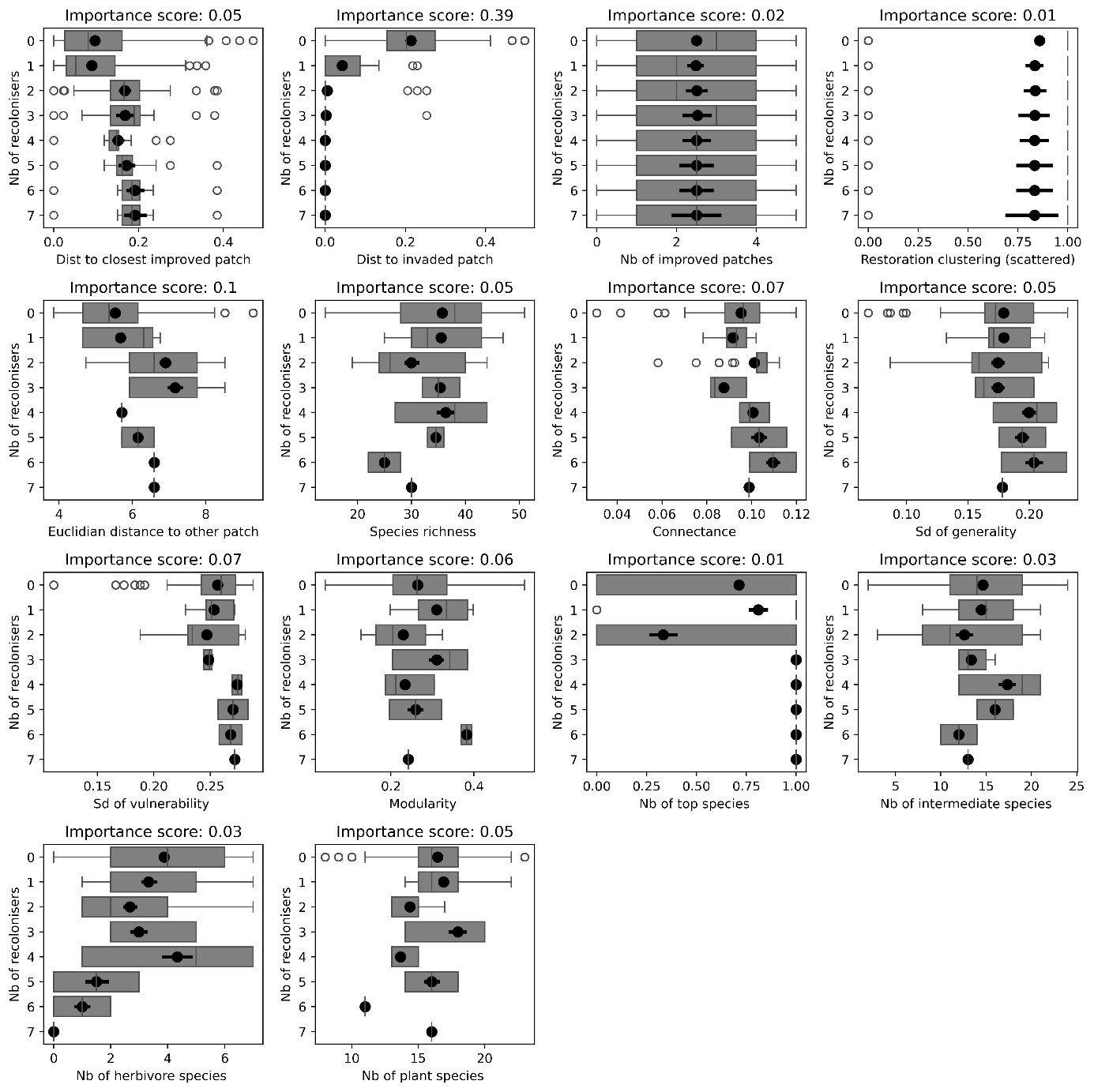

***Figure S1.2: Effects of food web topology, number of improved patch and location of recolonised patch on number of recolonisations from trophic level 0 (plants) across simulations with 0 to 5 patches restored.*** Boxplots represent median and 25% and 75% quantiles (quartiles). Black points and error bars represent the mean and 95% confidence interval around the mean. Each boxplot illustrates one investigated driver of recolonisation: from patch properties such as distance to improved patch and source patch, summed Euclidian distance to other patches (a measure of patch isolation), landscape properties such as the number of improved patches and restoration type (scattered = 1, clustered = 0), and properties of the food web before restoration (patch-level): species richness, connectance, standard deviation (SD) of generality, vulnerability, modularity and number of species per trophic level.

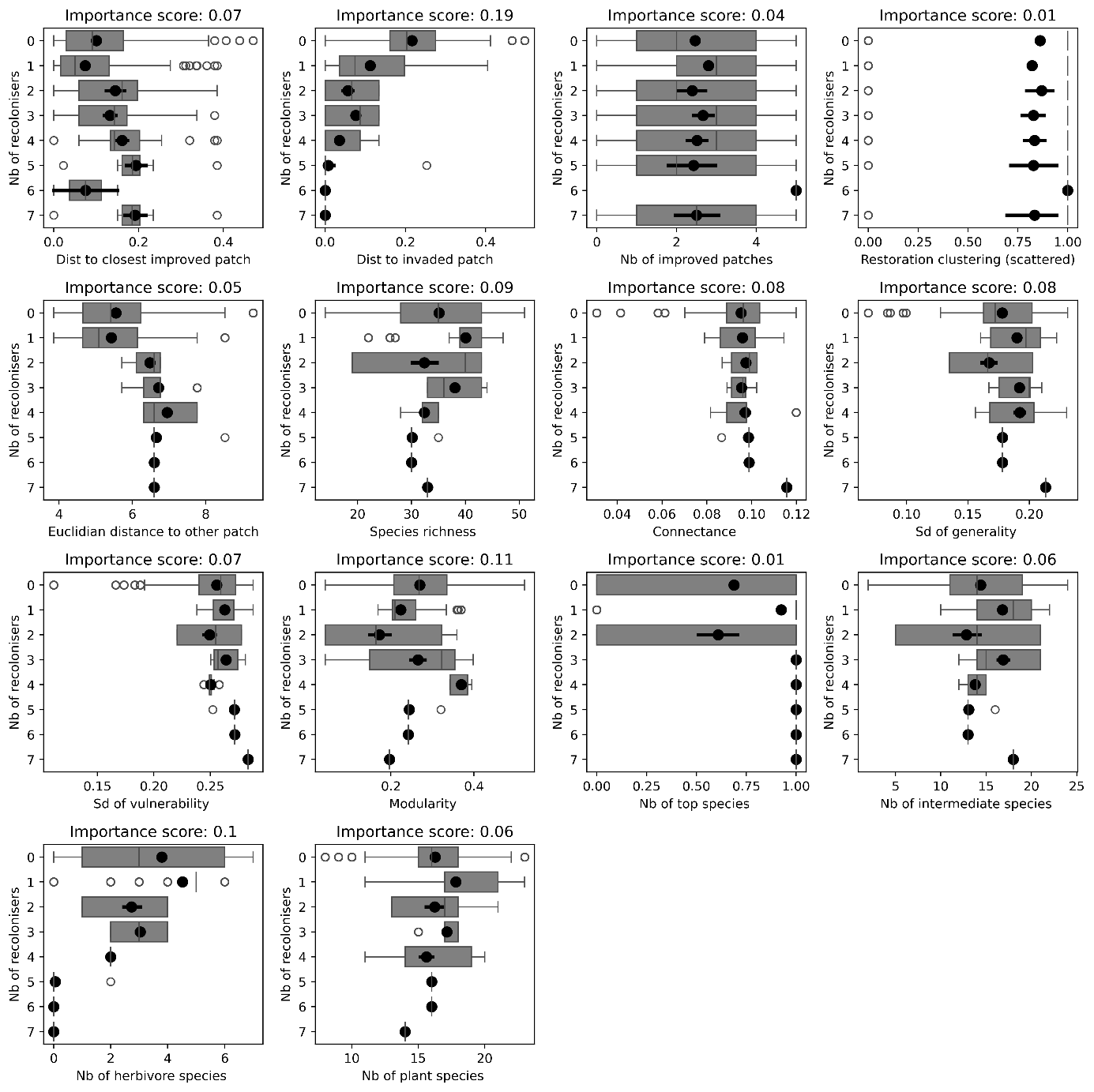

***Figure S1.3: Effects of food web topology, number of improved patch and location of recolonised patch on number of recolonisations from trophic level 1 (herbivores) across simulations with 0 to 5 patches restored.*** Boxplots represent median and 25% and 75% quantiles (quartiles). Black points and error bars represent the mean and 95% confidence interval around the mean. Each boxplot illustrates one investigated driver of recolonisation: from patch properties such as distance to improved patch and source patch, summed Euclidian distance to other patches (a measure of patch isolation), landscape properties such as the number of improved patches and restoration type (scattered = 1, clustered = 0), and properties of the food web before restoration (patch-level): species richness, connectance, standard deviation (SD) of generality, vulnerability, modularity and number of species per trophic level.

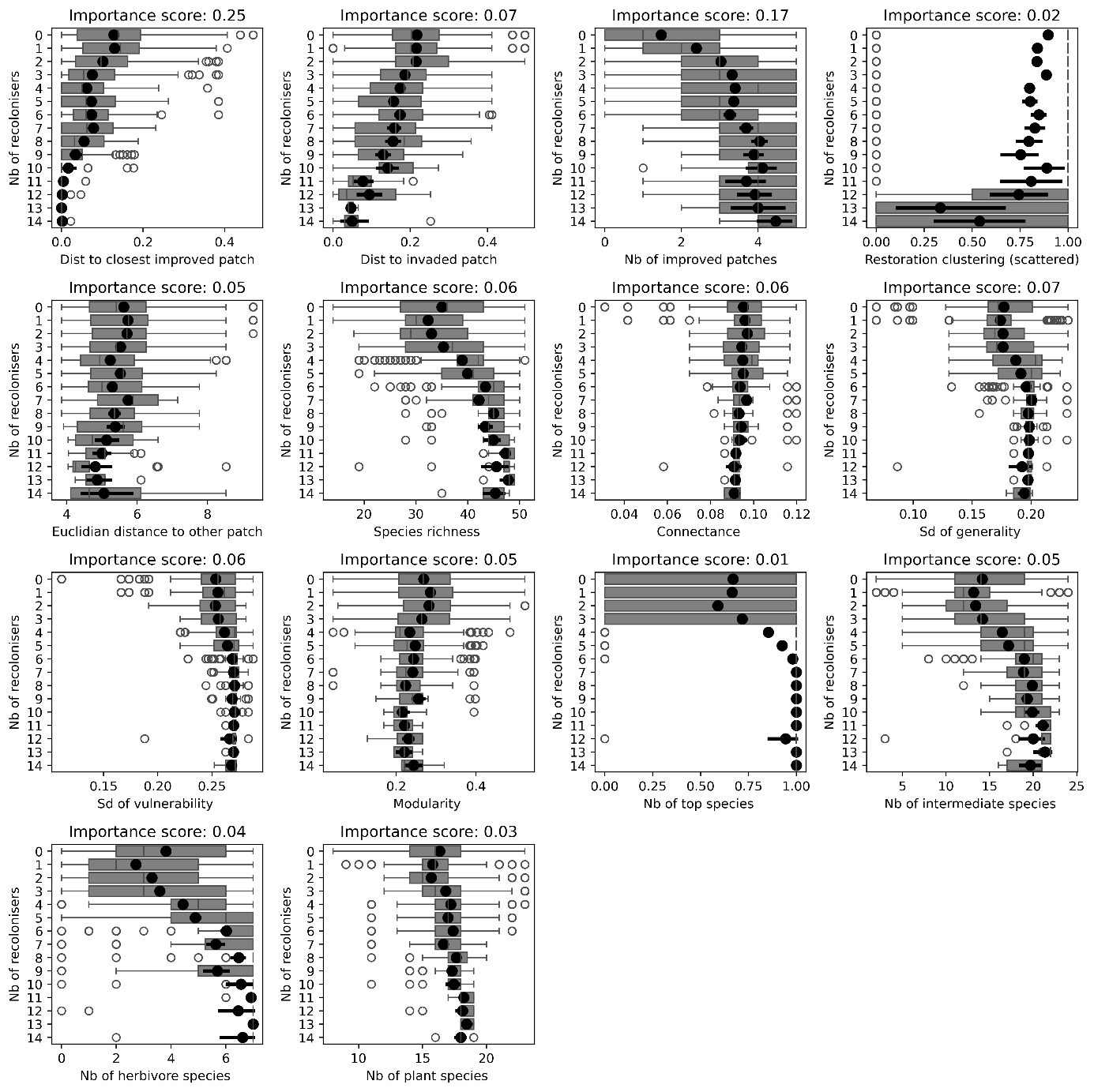

***Figure S1.4:*** ***Effects of food web topology, number of improved patch and location of recolonised patch on number of recolonisations from trophic level 2 (intermediate species) across simulations with 0 to 5 patches restored.*** Boxplots represent median and 25% and 75% quantiles (quartiles). Black points and error bars represent the mean and 95% confidence interval around the mean. Each boxplot illustrates one investigated driver of recolonisation: from patch properties such as distance to improved patch and source patch, summed Euclidian distance to other patches (a measure of patch isolation), landscape properties such as the number of improved patches and restoration type (scattered = 1, clustered = 0), and properties of the food web before restoration (patch-level): species richness, connectance, standard deviation (SD) of generality, vulnerability, modularity and number of species per trophic level.

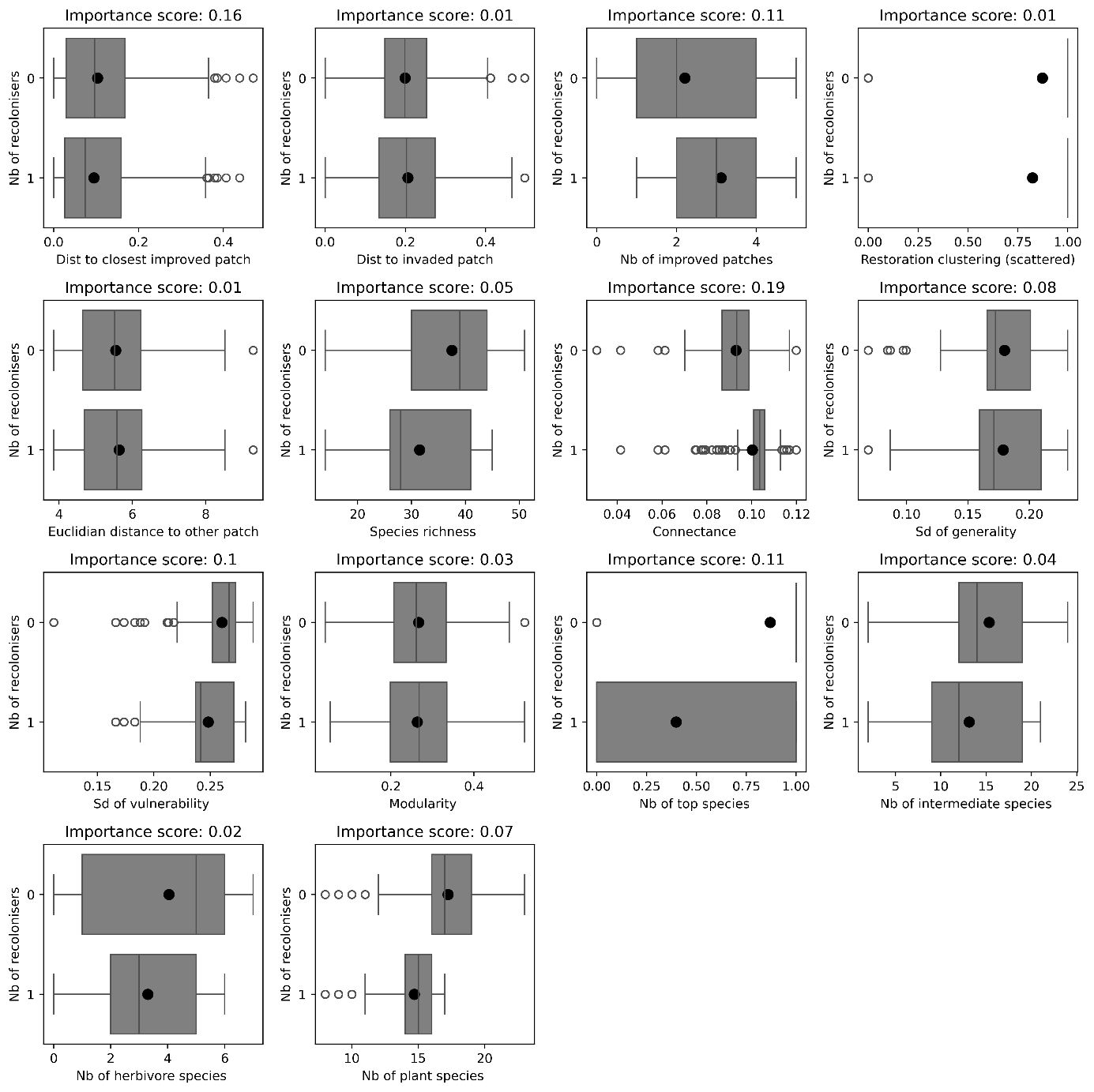

***Figure S1.5:*** ***Effects of food web topology, number of improved patch and location of recolonised patch on number of recolonisations from trophic level 3 (top species) across simulations with 0 to 5 patches restored*.** Boxplots represent median and 25% and 75% quantiles (quartiles). Black points and error bars represent the mean and 95% confidence interval around the mean. Each boxplot illustrates one investigated driver of recolonisation: from patch properties such as distance to improved patch and source patch, summed Euclidian distance to other patches (a measure of patch isolation), landscape properties such as the number of improved patches and restoration type (scattered = 1, clustered = 0), and properties of the food web before restoration (patch-level): species richness, connectance, standard deviation (SD) of generality, vulnerability, modularity and number of species per trophic level.

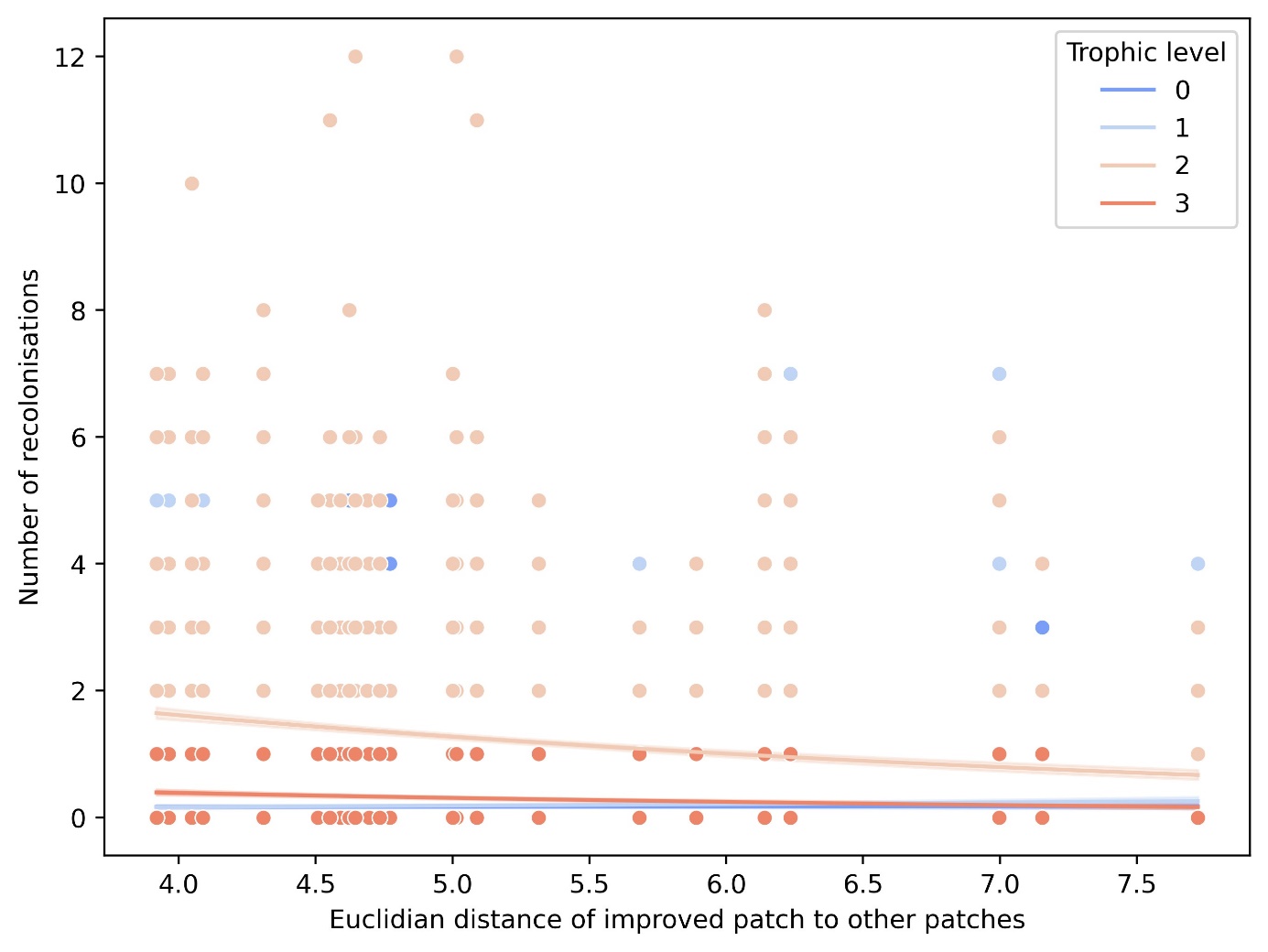

***Figure S1.6: Effect of patch-connectivity of the first improved patch on the number of recolonisations across trophic levels and all patches.*** Regression lines are fitted from a poisson generalised linear model of the relationship between number of recolonisers and the interaction between Euclidian distance of improved patch to other patches (patch isolation) and trophic level. Points represent number of recolonisations at the local (patch) level across all patches for the simulations where only one patch was restored.

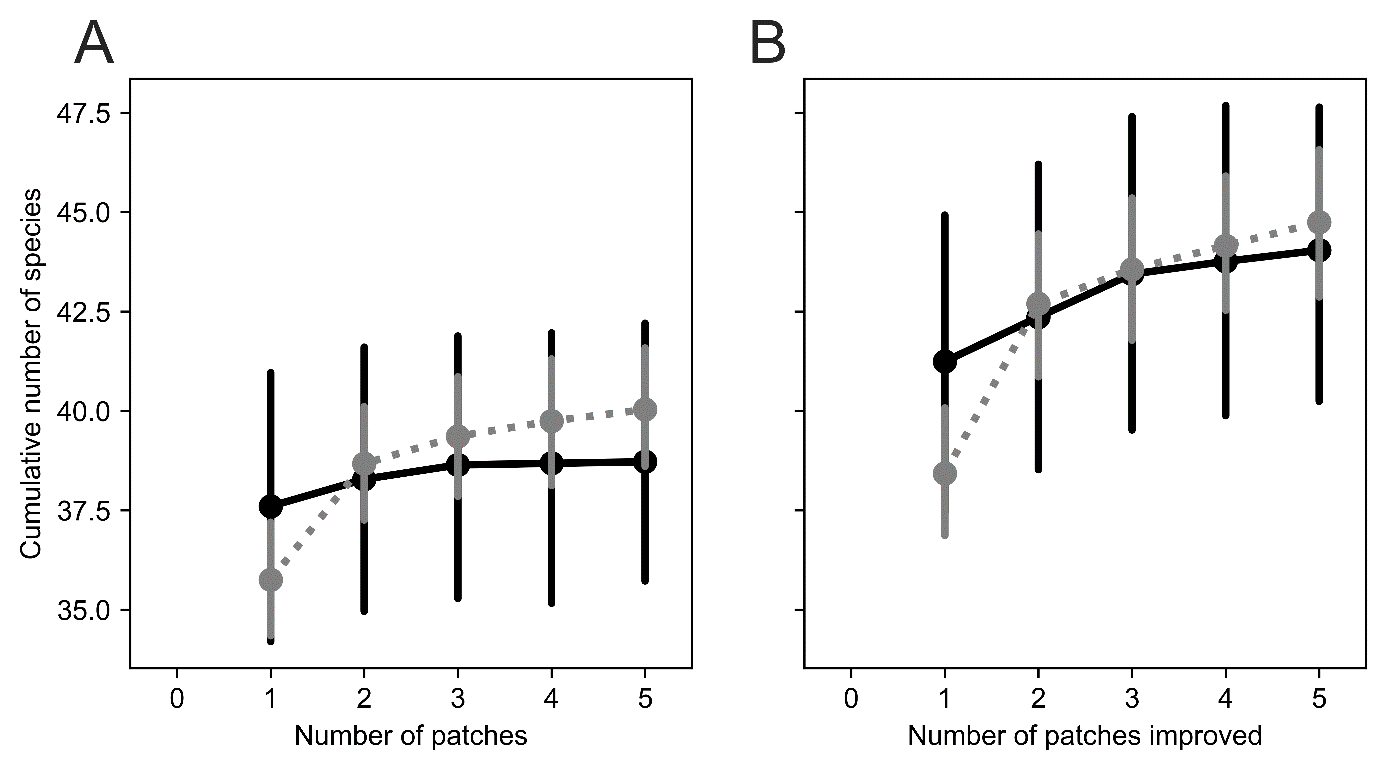

**Figure S1.7: No strong effect of clustering on restoration outcome.** Species accumulation curves (SACs) are presented for to-be-restored patches in initial simulations, before restoration (A) and for restored patches after restoration (B) for clustered and scattered scenarios: clustered (black continuous line) where the five improved patches were clustered in the middle of the landscape or scattered (grey dotted line) scenario where improved patches were randomly scattered in space.

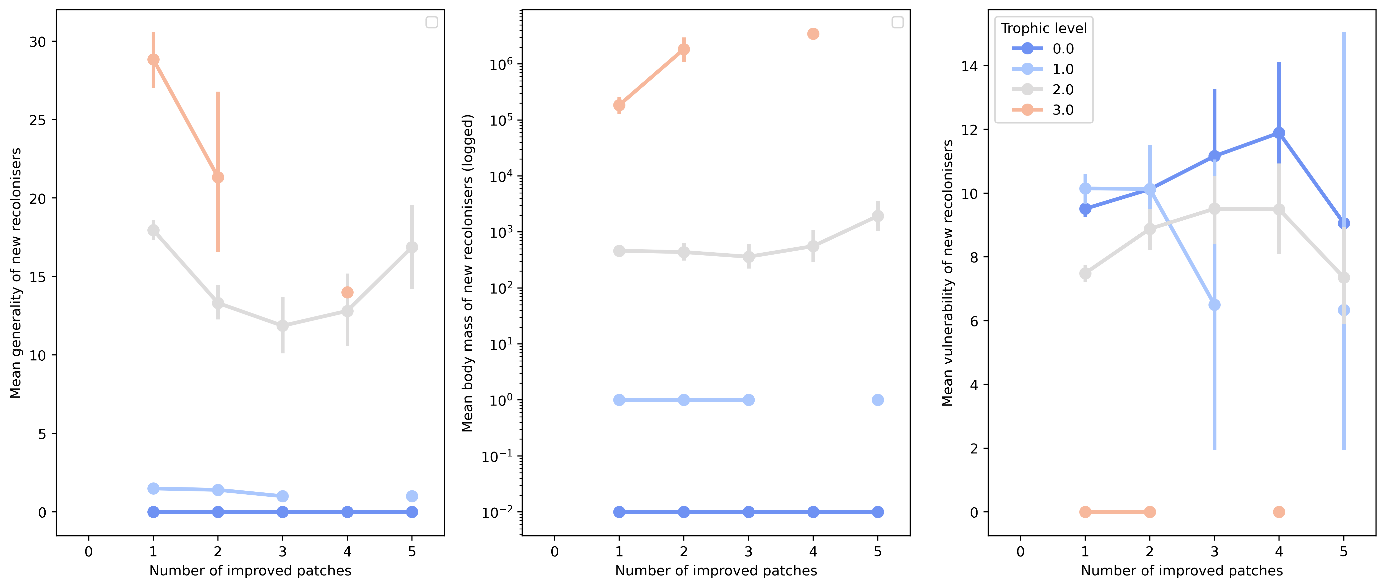

**Figure S1.8: Mean traits of new recolonisers across trophic levels on restored patches only.** Each plot shows the evolution of mean traits (generality, body mass and vulnerability) of new invaders across trophic levels as more patches are improved. Points represent the mean trait of new recolonisers and error bars the 95% confidence interval around the mean. Missing points mean that no recolonisation event were ever recorded for that improvement level x trophic level. Generality of top predators seems to decrease with further patch improvement, but more recolonisation events would be needed to confirm this trend.
